# Selection-like biases emerge in population models with recurrent jackpot events

**DOI:** 10.1101/182519

**Authors:** Oskar Hallatschek

## Abstract

Evolutionary dynamics driven out of equilibrium by growth, expansion or adaptation often generate a characteristically skewed distribution of descendant numbers: The earliest, the most advanced or the fittest ancestors have exceptionally large number of descendants, which Luria and Delbrück called “jackpot” events. Here, we show that recurrent jackpot events generate a deterministic bias favoring majority alleles, which is equivalent to an effective frequency-dependent selection (proportional to the log ratio of the frequencies of mutant and wild-type alleles). This “fictitious” selection force results from the fact that majority alleles tend to sample deeper into the tail of the descendant distribution. The flipside of this sampling effect is the rare occurrence of large frequency hikes in favor of minority alleles, which ensures that the allele frequency dynamics remains neutral overall unless genuine selection is present. The limiting allele frequency process is dual to the Bolthausen-Sznitman coalescent and has a particularly simple representation in terms of the logarithm of the mutant frequency. The resulting picture of a selection-like bias compensated by rare big jumps allows for an intuitive understanding of allele frequency trajectories and enables the exact calculation of transition densities for a range of important scenarios, including population size changes and different forms of selection. The fixation of unconditionally beneficial mutations is shown to be exponentially suppressed and balancing selection can maintain diversity only if the population size is large enough. We briefly discuss analogous effects in disordered complex systems, where sampling-induced biases can be viewed as ergodicity breaking driving forces.

One of the virtues of mathematizing Darwin’s theory of evolution is that one obtains quantitative predictions about the dynamics of allele frequencies that can be tested with increasing rigor as experimental techniques, sequencing methods and computational power advance. The Wright-Fisher model is arguably the simplest null model of how allele frequencies change across time [17]. Although, for modeling neutral genetic diversity, it is often replaced by equivalent backward-in-time models of the ensuing tree structures [23], forward-in-time approaches are still unrivaled in their ability to include the effects of natural selection. As such, the Wright-Fisher model has been instrumental for shaping the intuition of generations of population genetics about the basic dynamics of neutral and selected variants. But transition densities derived from the Wright-Fisher model also find tangible application in scans for selection in time series data [4, 9, 22].

The Wright-Fisher model is remarkably versatile as it can be adjusted to many scenarios by the use of *effective* model parameters: An effective population size, an effective mutation rate and effective selection coefficients. But, crucially, these re-parameterizations cannot account for extremely skewed family size distributions. While remarkably skewed family distributions occur in some natural populations [15], they routinely arise in microbial populations that combine exponential growth with recurrent mutations. This was first highlighted by Luria and Delbrück [26], who noticed that mutations that occur early in an exponential growth process will produce an exceptionally large number of descendants. The distribution of such mutational “jackpot” events has a particular power law tail in well-mixed population, as is briefly explained in Fig. 1A. Simplest models of continual evolution [28] and related models of traveling waves [36] can be viewed, on a coarse-grained level, as repeatedly sampling from this jackpot distribution. (The number of draws and the characteristic resampling time scale varies with the model.) It is by now well-established that the ensuing genealogies are described by a particular multiple-merger coalescent [7, 8, 13, 29, 33–35] first identified by Bolthausen and Sznitman [5].

**FIG. 1:**
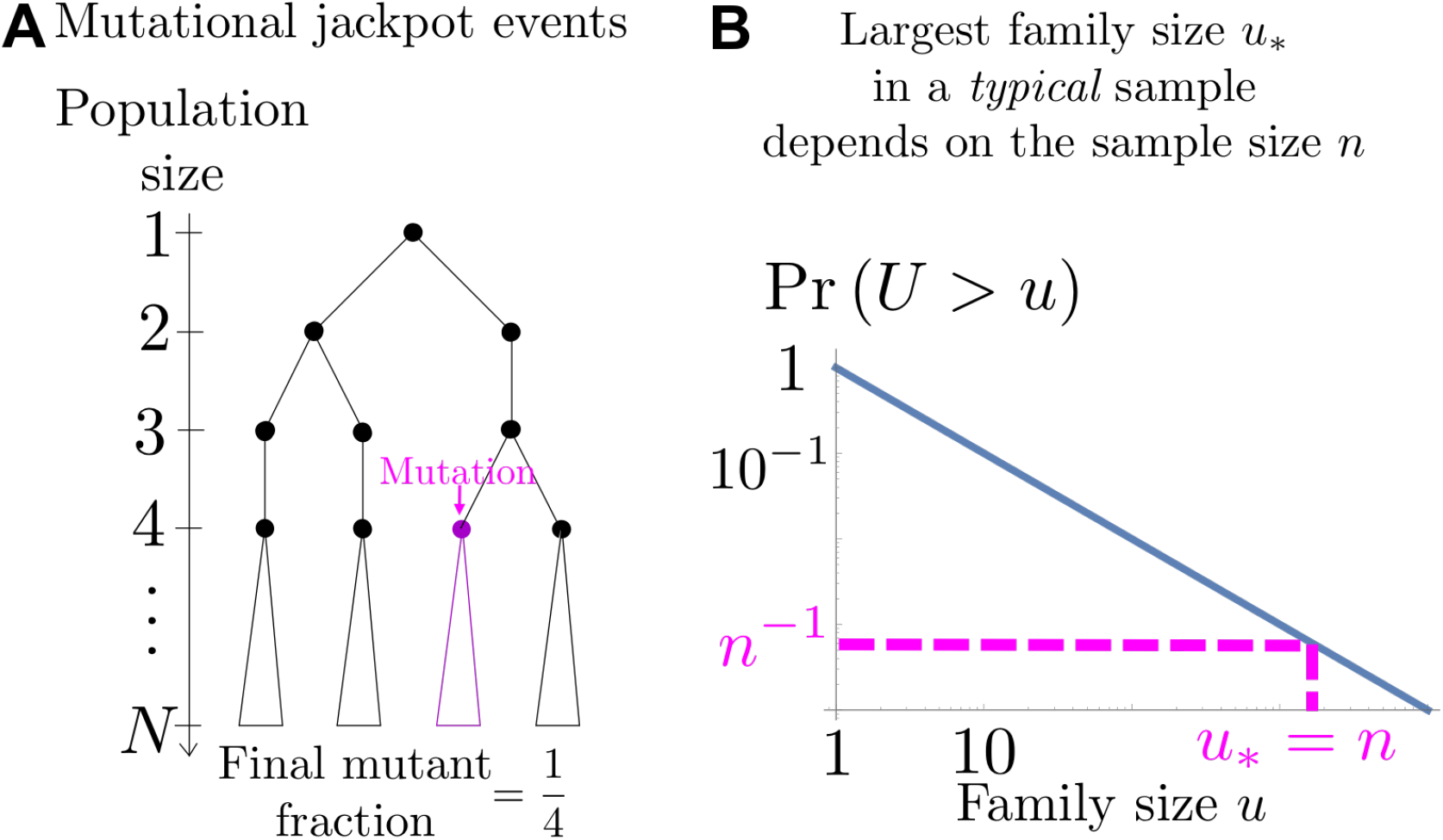
A) Illustration of mutational jackpot events, first studied by Luria and Delbriick. Consider the growth of a well-mixed population of microbes starting from a single cell (ignore death). A mutation that occurs in the j^th^ cell division will have a final frequency of about ≈ *1/j* (j = 4 in the illustration) and thus a “family size” of *u* = *N/j*. Hence, the probability Pr[*U* > *u*] to reach an even larger family size *U* is equal to the probability ≈ *j/N* = 1/u that the mutation occurs prior to the n^th^ cell division. The probability density to acquire a family size *u* therefore exhibits a power law tail *p*(*u*) ∝ *u*^-2^. (Our argument ignores the stochasticity in cell division events, which however does not change the power law exponent.) B) The blue line indicates the probability Pr[*U* > *u*] that the family size *U* of a mutation is larger than *u*. The largest family size *u*_*_ in a sample of *N* jackpot events should typically be of order *N* because the probability of sampling an even larger event is 1/n (dashed lines). A typical n-sample, therefore, has a mean family size of order log(*n*), which is obtained upon truncating the family size distribution at *u*_*_. It turns out that this effect generates a selection-like bias favoring majority alleles, which is compensated by rare sampling events that favor minority alleles.

While extensions of the Wright-Fisher diffusion process to capture skewed offspring numbers have been formally constructed [2, 3, 14, 20], also including selection and mutations [1, 10, 11, 16, 18], we still lack explicit finite time predictions for the probability distribution of allele frequency trajectories. Our goal here is to fill this gap for the particular case of the Luria-Delbriick jackpot distribution, by characterizing the allele frequency process in such a way that it can be easily generalized, intuitively understood and integrated in time.

## I. SAMPLING ALLELE FREQUENCIES ACROSS GENERATIONS

Our starting point is a simple model for the dynamics of a subpopulation at frequency *X*(*t*), the mutants, within a population of total size *N*(*t*) passing through discrete generations *t* = 1, 2,… Note that the population size is allowed to change from generation to generation. The mutant frequency *X*(*t* +1) in the generation *t* + 1 is produced from generation *t* in two steps: First, each individual gets to draw a statistical weight *U* from a given probability density function *p*(*u*) (nonzero only for *u >* 0 and the same for both mutants and wild-types). This generates *NX*(*t*) random variables 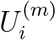 for the mutants and *N*(1 — *X*(*t*)) random variables 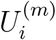 for the wild-types. The new discrete mutant number *NX*(*t* + 1) is obtained in a second step by binomially sampling N times with the success probability

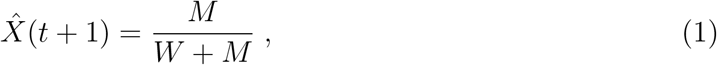

which depends on the sums 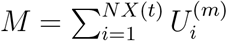 and 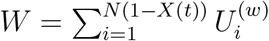. One can think of the statistical weights *M* and *W* as representing the total number of mutant and wild-type “seeds” in a large seed pool, from which only a finite number *N* (sampled with replacement) go on to survive to adulthood. Note that the deviation of the new mutant frequency *X*(*t* + 1) from 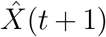, the simple fraction Eq. 1, is of order 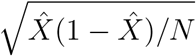. For most of this work, we are interested in large enough *N* and a broad enough descendant distribution such that the binomial sampling error, which represents classical random genetic drift, is negligible compared to the fluctuations induced by sampling from the descendant distribution. Also note that because all random variables are independently drawn from the same distribution, there is no expected bias in the mutant frequency: The expected allele frequency in generation *t* is constant and equal to the starting allele frequency (*X*(*t*) is a “martingale”).

If *p*(*u*) has finite mean and variance, binomial sampling does matter and the above population model leads to Wright-Fisher diffusion in the large *N* limit [30], which has genealogies described by the Kingman coalescent [33].

But what are the characteristic features of the forward allele frequency process *X*(*t*) for descendant distributions so broad that even the mean diverges? As I show heuristically in the next section, this leads to a sampling-induced bias for alleles that are in the majority, an apparent rich-get-richer effect. I will then describe and illustrate phenomena driven by this effect, including biased time series, a high-frequency uptick in site frequency spectra and a low probability of fixation of beneficial mutations. In the technical section of this paper, I discuss a suitable large-*N* scaling limit of *X*(*t*), in which the stochastic dynamics can be fully predicted.

## II. TYPICAL OFFSPRING NUMBERS FAVOR THE MAJORITY TYPE

It is useful to first consider purely heuristic arguments to see that, for broad enough descendant number distributions, population resampling typically favors the majority type. These arguments provide an intuitive basis for the phenomena I discuss and derive further below.

Suppose an allele is currently at frequency *X*, and we would like to estimate the frequency *X’* after resampling. According to our reproduction rules stated above, we need to estimate the total number of descendants of both mutants, *M*, and wild-types, *W*, which represent sums of many random family sizes (if *N* is large). Such estimates are challenging for skewed family size distributions, especially when the mean depends on the largest families that occur in a sample. Yet, a mean family size ⟨*U*⟩_*n*_ of a *typical* sample of *N* offspring numbers {*U_i_*}_*i*=ı…*n*_ can be estimated by the truncated expectation [31]

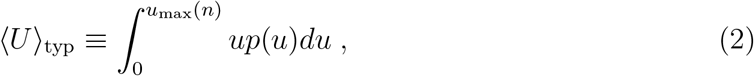

The cutoff *u*_max_ (*n*) of the integral represents the largest family size in a typical *n*-sample, illustrated in Fig. 1B, which can be estimated by the extremal criterion[25]

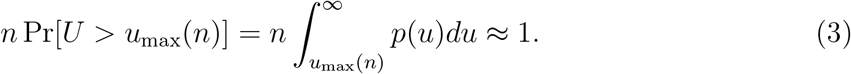

Note that the typical value ⟨*U*⟩_typ_ becomes essentially equal to the expectation ⟨*U*⟩ if the integral in Eq. 2 is not sensitive to the upper bound. The key phenomena discussed in the paper, however, rely on *p*(*u*) being sufficiently broad-tailed so that ⟨*U*⟩_typ_ is *dependent* on the sample size *n*. This occurs when *p*(*u*) decays like *u*^−2^ or more slowly, so that the mean family size diverges. Throughout most of this paper, I will in fact focus on the particularly interesting marginal case *p*(*u*) ~ *u*^−2^, which as mentioned arises in population models that combine stochastic jumps with exponential growth. In this case,

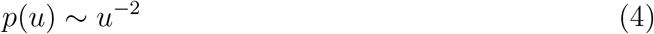

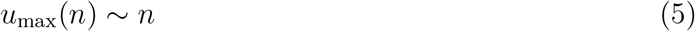

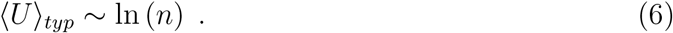

Hence, the family size of a typical sampled from a mutant population currently at frequency *X* can be estimated by

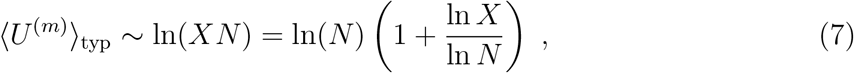

and likewise for the wild-type population. The frequency-dependence of this expression is the crux of our study, as it implies that more abundant mutants (larger *X*) typically behave as if they have higher fitness (larger family sizes).

Assuming an initial allele frequency *X*, the typical frequency (*X*′)_typ_ after resampling can then be estimated by

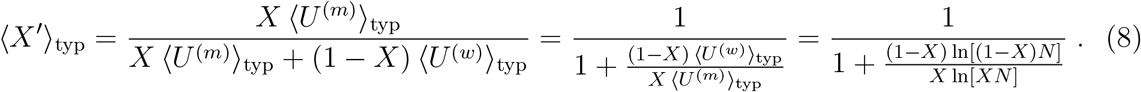

The typical discrepancy between the resampled frequency (*X*′)_typ_ from the original frequency *X* simplifies in the limit | ln N| ≫ | ln *X* | to a deterministic advection velocity

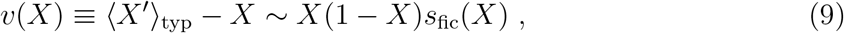

which has the form as a traditional selection term with a frequency-dependent selection coefficient

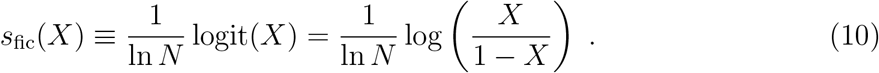

The selection-like advection velocity tends to increase the frequency of the majority type and results from the fact that the majority type is able to sample deeper into the tail of the offspring number distribution. I will call this effect *fictitious* selection because it acts like selection, yet is of purely probabilistic origin. Both the advection velocity and the fictitious selection coefficient are plotted in Fig. 2.

**FIG. 2:**
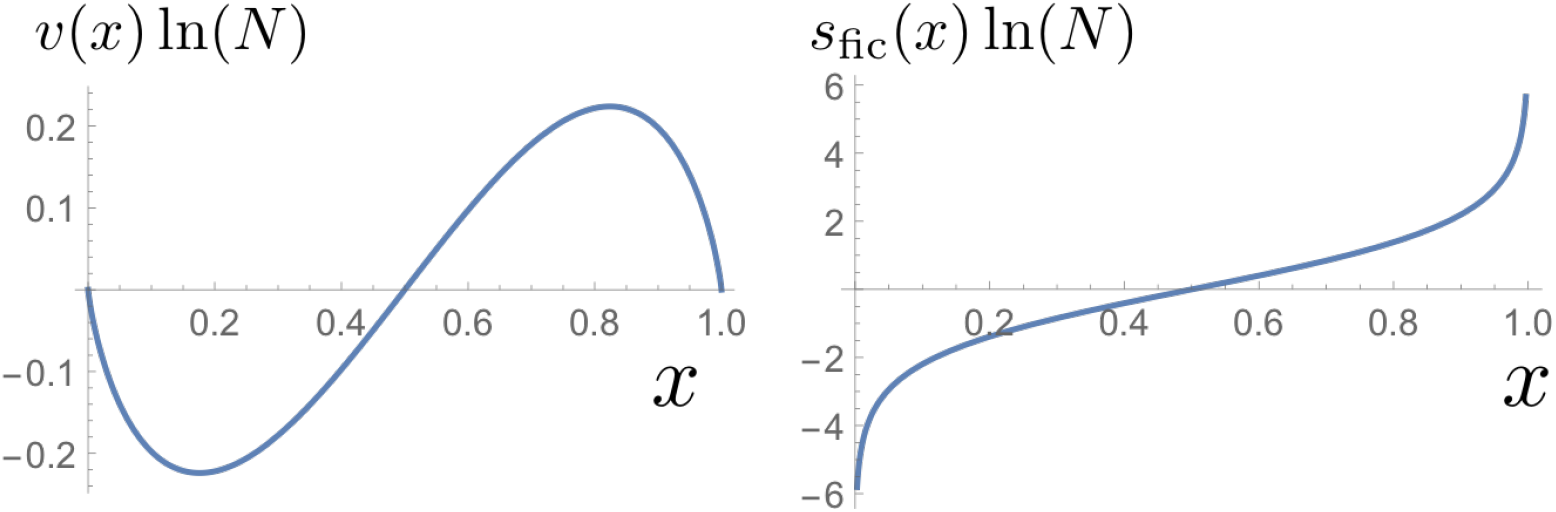
Scaled advection velocity *v*(*x*) and fictitious selective difference *s*_fic_(*x*) as a function of mutant fraction *x*. The advection tends to push the frequencies of polymorphic sites towards fixation and extinction, depending on what boundary is closer. The bias slows down near the boundaries, which leads to an accumulation of high and low frequency variants in site frequency spectra.

One naturally wonders how the fictitious selection force can be consistent with the overall neutrality of the process: The average mutant frequency has to stay constant in time. But, so far, we have only considered the typical behavior of the allele frequency dynamics, and it turns out that the ignored atypical events ensure neutrality: A detailed analysis of the resampling distribution (SI Sec. X) shows that neutrality holds overall because rare big events rescue the minority type. Both effects, a nearly deterministic advection towards the majority type and compensating rare jumps in favor of the minority type, can be appreciated from histograms that show resampled frequencies conditional on an initial frequency *X*_0_ ≪ 1. For large *N*, a pronounced peak appears below *X*_0_, at a scale consistent with the selection coefficient determined above, see Fig. 3. Yet, the histogram still has some support at very large frequencies, which correspond to the just-mentioned compensating jumps.

**FIG. 3:**
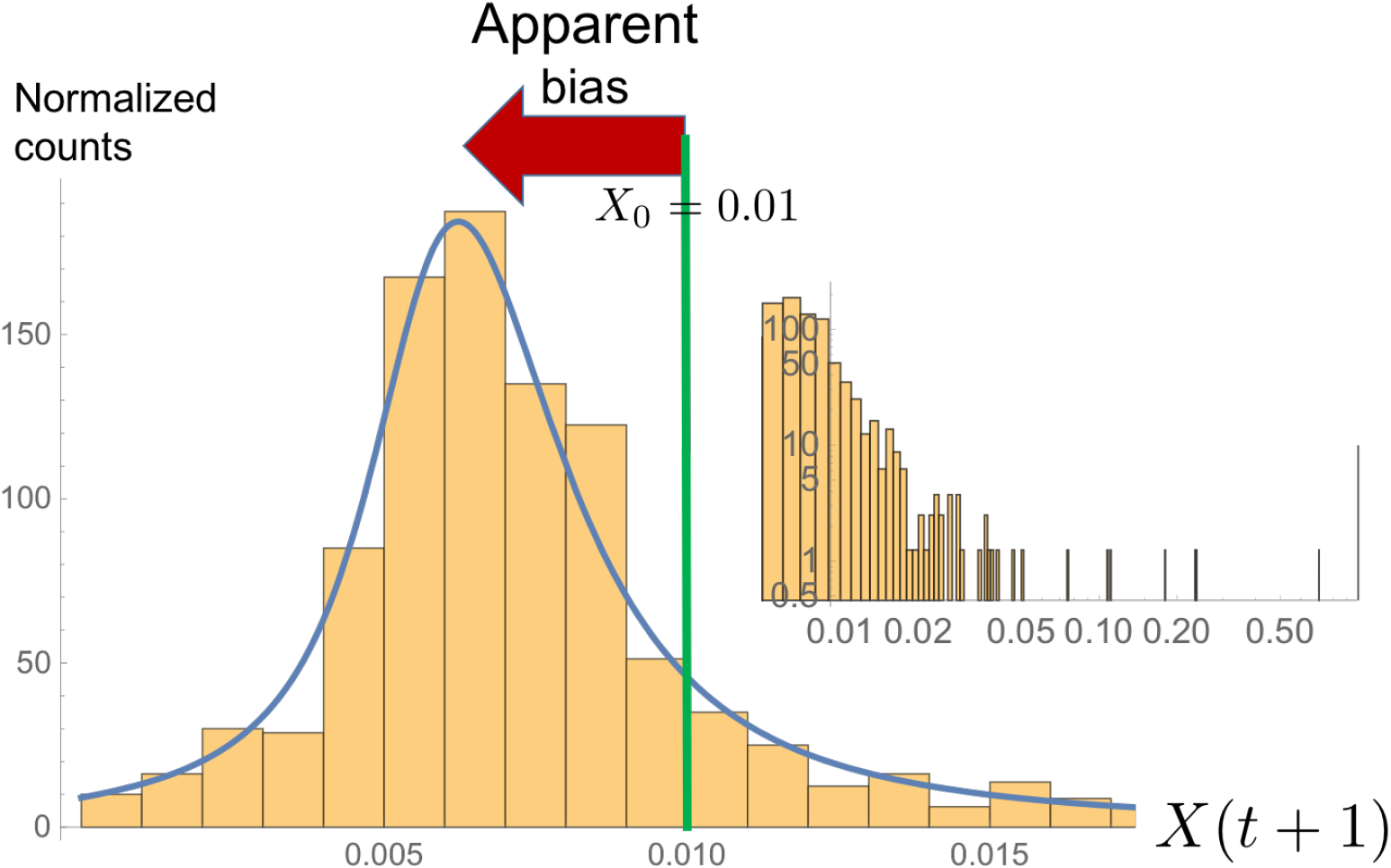
Histogram obtained from resampling 800 times a new allele frequency *X*(*t* + 1) given that *X*(*t*) = *X*_0_ = 0.01. The population size is *N* = 10^6^. Left: Notice the apparent shift to lower frequencies of the bulk of the histogram compared to the initial frequency (green line). The blue line shows the asymptotic resampling distribution (Eq. 79). Inset: Rare big events rescue neutrality. The top five events are {0.71, 0.24, 0.24, 0.18, 0.11}.

**FIG. 4:**
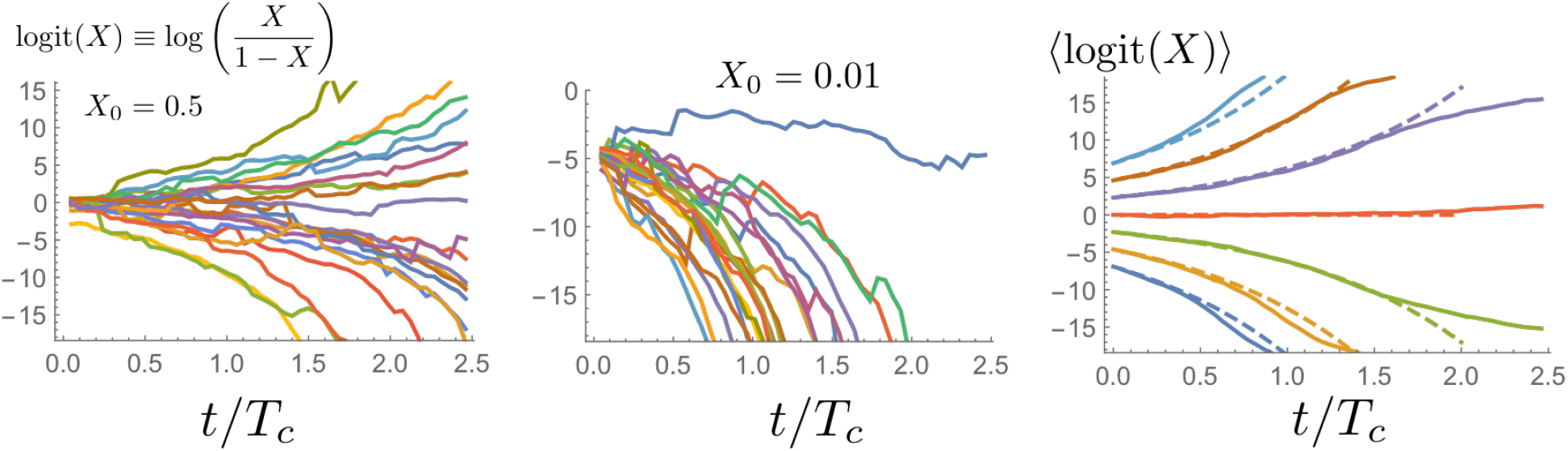
Sample paths for the log ratio of mutant frequency and wild-type frequency. The population size was *N* = 10^9^, which sets the coalescence time *T*_*c*_ = ln(*N*), and trajectories were chosen to start at *X*_0_ = 0.5 in the left panel and *X*_0_ = 0.01 in the middle panel, respectively. The panel on the right shows the behavior of the log ratio (solid lines) for different starting frequencies (averaged over 100 sample paths). The dashed lines are the theoretical expectation, which is an exponential in logit space (Eq. 12). **Also show trajectories in frequency space**.

## III. CONSEQUENCES

I will now describe tangible phenomena driven by fictitious selection and the compensating rare jumps. The mathematical description of these phenomena follows from an analytically solvable mathematical framework described in Sec. IV below.

### A. Trajectories

If we ignore the effect of rare jumps, one would expect a characteristic allele frequencies trajectories to move towards fixation or extinction, following the fictitious selection force. The resulting deterministic trajectory is most easily described in terms of the logit, Ψ(*t*) = logit[*X*(*t*)] = log[*X*/(1 − *X*)], which obeys

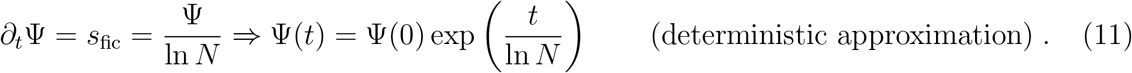

It turns out that the expectation of the logit indeed follows these dynamics,

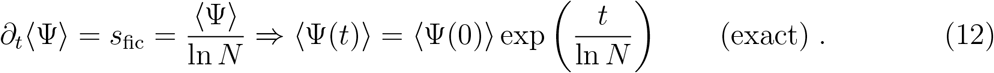

as the simulations in Fig. 6 demonstrate. Thus, while neutrality requires the expectation of allele frequencies to remain constant, one finds that the expectation of the log ratio of frequencies moves away from 0 and, more over, exponentially fast. This peculiar behavior reflects the fact that a log-transformed stochastic variable is much less sensitive to rare, big jumps, which are needed to compensate for the fictitious selection force. In frequency-space, the trajectories resemble deterministic trajectory pieces glued together by rare compensating jumps, as can be seen in the trajectories in Fig. 6.

The dynamics of the logit expectation in Eq. 12 implies that it typically takes a time of order ln *N* generations to change the allele frequency by order 1 and ln *N* ln ln *N* generations to reach one of the absorbing boundaries, which have a logit frequency of order *O*(ln(*N*)). Both time scales are known from the associated Bolthausen-Sznitman coalescent: *T*_c_ ≡ ln *N* is the time to coalesce a random sample of two lineages and *T*_*c*_ ln ln *N* is the time scale to coalesce *all* lineages in the population [2].

Fictitious selection can be most easily detected at the high-frequency end of the site frequency spectrum (SFS). As with regular selection, the SFS near fixation solely depends on the advection term, SFS(*x*) ~ 1/*v*(*x*) as *x→* 1, as if allele frequencies are only advected and jumps are negligible. This leads to an uptick at high frequencies [29] that has the same form as the SFS of a selected allele with frequency-dependent selection coefficient *s*_fic_(*x*). The excess of common alleles relative to intermediate frequencies results from the advection term slowing down as allele frequencies increase towards fixation.

It turns out that the joint action of fictitious selection and compensating jumps can be best represented in logit space. The logit of the frequency, Ψ(*t*) ≡ logit(*X*(*t*)), continuously switches between exponentially diverging deterministic trajectories via jumps drawn from a jump kernel, which is a power law for small jump distances but exponentially decaying for large jumps. The ensuing stochastic process is illustrated in Fig. 5.

**FIG. 5:**
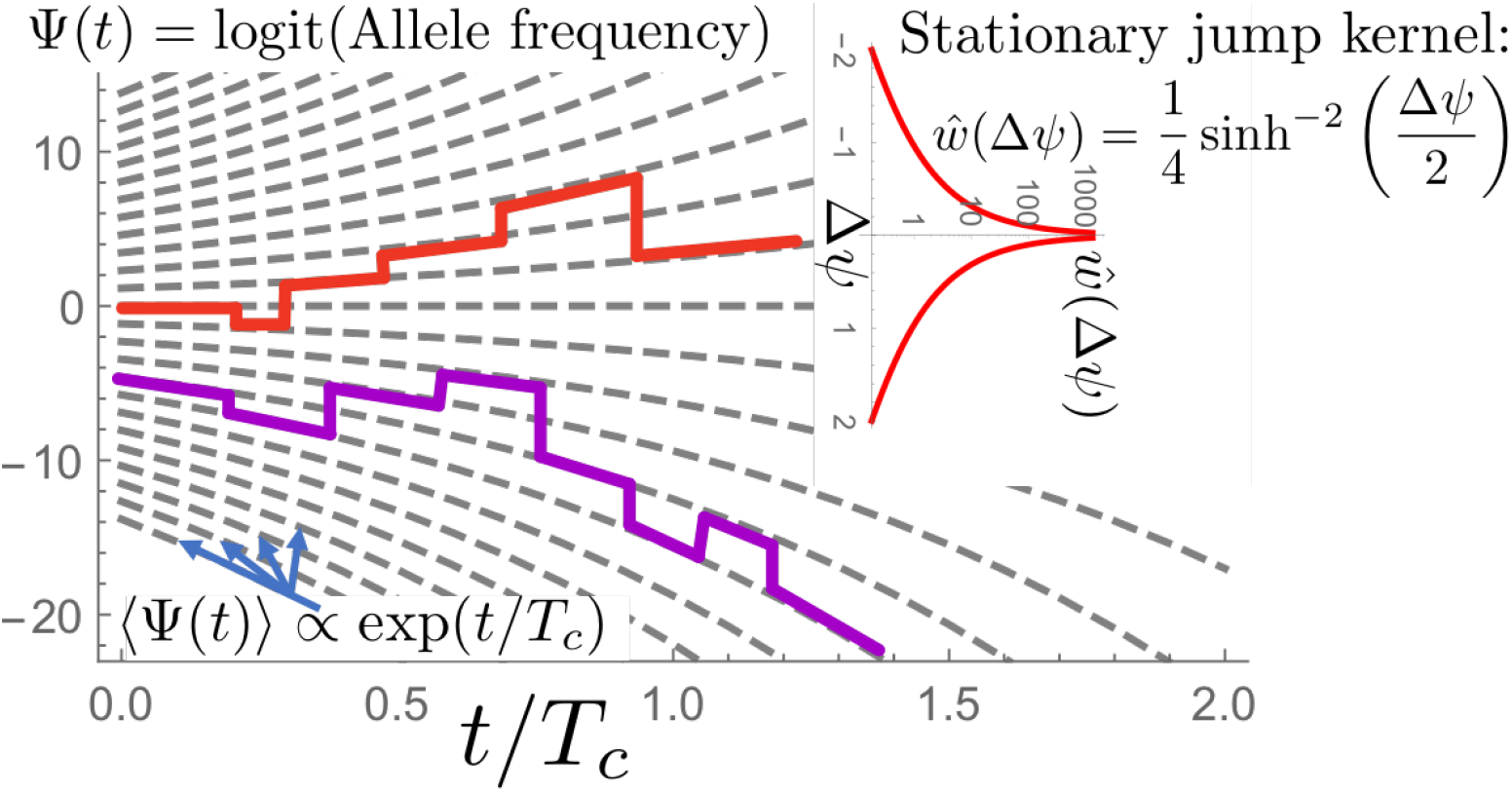
Illustration of the limiting allele frequency process, which is most easily described for the log-ratio of the frequency of mutants and wild-types, Ψ(*t*) = logit[*X*(*t*)] ≡ log(*X*/(1 − *X*)). Trajectories in this logit space on average follow exponentially diverging paths (gray lines). Switching between these paths occurs at a rate 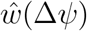, which depends (only) on the distance △*ψ* in logit space. Small jumps happen frequently, in fact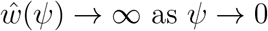, but impactful jumps of order 1 or larger roughly take a coalescence time *T*_*c*_ = ln *N* to occur. The advection force drives the ultimate fixation or extinction of alleles, which takes a time of order *T*_*c*_ ln ln *N*.

Interestingly, the jump rate between two positions in logit space only depends on their distance (the kernel is stationary). This makes it possible to compute exactly for a general time-dependent population size the probability density 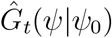 that a trajectory will move from logit position *ψ*_0_ to *ψ* in a time period *t*. This transition density is given by

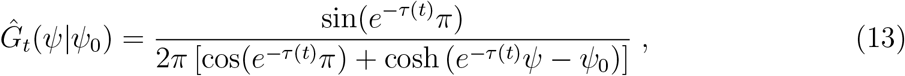

in terms of a rescaled time variable 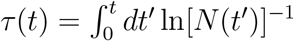, which simplifies to *τ* = *t* ln(*N*)^−1^ in the case of a constant population size. The scale factor ln[*N*(*t*)]^−1^ appearing in this time conversion represents the coalescence rate of two lineages at time *t*. For a constant population size, the time conversion simplifies to *τ* = *t*/*T*_*c*_ where *T*_*c*_ is the mean coalescence time of two lineages. Using the transition probability in Eq. 33 it is possible to characterize a number of interesting statistics, such as sojourn times or the site frequency spectrum, and directly confirm a duality with the Bolthausen-Sznitman coalescent.

### B. Selection

Selection modifies the above dynamics by introducing a true bias (no jumps to leading order). A mutation with frequency-independent fitness effect s leads to

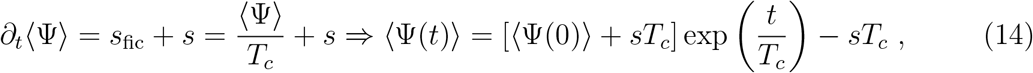

where a constant coalescence rate 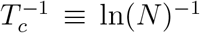 is assumed. From this expression, we can see that genuine selection indeed competes with fictitious selection: The logit will on average increase with time only if ⟨Ψ(0)⟩ + s*T*_c_ is larger than 0, otherwise it will decay over time. Mean sample paths are shown in

In fact, our detailed analysis below will show that the stochastic dynamics of a selected allele starting at frequency *ψ*(0) is identical to the dynamics of a neutral allele properly shifted in logit space. The exact statement is

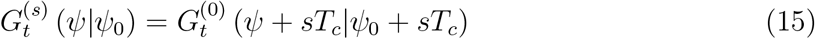

in terms of a transition density 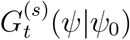 of a selected allele with selective advantage s. Since the fixation probability of a neutral allele at frequency *X* simply is *X*, this mapping implies

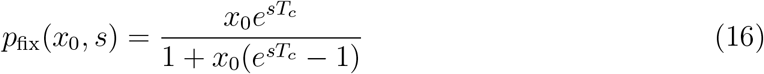

If the selected allele is initially rare, *x*_0_ ≪ 1, we have 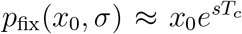. So, a selective advantage does increase the odds of fixation exponentially. But, because *x*_0_ = *O*(*N*^−1^) for a single mutant, the fixation probability of newly arising beneficial mutations is very small, unless *s*T*_*c*_* = *O*(ln*N*).

Finally, since the effective bias tries to push allele frequencies towards fixation, one may ask what happens in presence of balancing selection opposing fictitious selection. So, let us consider an allele under balancing selection modeled by a s(*ψ*) = — a(*ψ* — *ψ*_c_) in the sense of a Taylor expansion in logit space. The first moment in logit space now satisfies

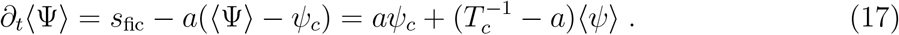

The most important feature of this expression is a threshold phenomenon at *a_c_* = **T*_*c*_*^−1^. For *a* < *a*_*c*_, balancing selection will not be able to maintain diversity. For *a* > *a*_*c*_ both alleles will be maintained in a balancing selection-draft equilibrium, with a wide variance if a is close to *a*_*c*_.

## IV. LIMITING STOCHASTIC PROCESS

I now show how the above phenomena follow from a detailed mathematical analysis of the asymptotic allele frequency dynamics as the population size tends to infinity. The resulting process belongs to the class of Lambda-Fleming-Viot processes, which are dual to multiple merger coalescents in a similar way as Wright-Fisher diffusion is dual to the Kingman coalescent. Although Lambda-Fleming-Viot processes [2, 3, 14] and extensions involving selection and mutations [1, 10, 16, 18] have been extensively studied, closed-form predictions for the transition density of allele frequency trajectories are still lacking. It turns out that, upon formulating the process in terms of a jump-drift process, such closed-form predictions can be obtained for the particular case of the Luria-Delbriick family size distribution, which generates a Lambda-Fleming-Viot process dual to the Bolthausen-Sznitman coalescent. Although somewhat technical in nature, the analysis will elucidate how fluctuations, the sampling-induced bias and an actual bias combine to control the fate of alleles, pointing to further theoretical directions that could be explored. Yet, readers not interested in mathematical details may jump right to the Discussion section.

The larger the population size *N* the more deterministic is the resampling of allele frequencies, and the slower the ensuing stochastic process. Hence, to obtain an interesting time-continuous stochastic process, it is natural to slow down the progression of time. It turns out that a well-behaved, time-continuous, Markov process *X*(τ) is obtained in terms of the time variable *τ* with differential *dτ* = *dt*/log[*N*(*t*)] upon sending log(*N*) → ∞. Since 1/log[*N*(*t*)] proves to be the coalescence rate for two lineages, one can say that we measure time in units of the inverse coalescence rate. For a constant population size, one simply has *τ* = *t*/log(*N*) = *t*/*T*_c_, where *T*_c_ is the mean coalescence time of two lineages.

Any such time-continuous (sufficiently well-behaved) Markov process is defined by an advection velocity, diffusion coefficient and jump kernel [19]. To state this triplet, we define *w_N_*(*x*_2_|*x*_1_) to be the probability density to sample a frequency *x*_2_ if we start with a frequency *x*_1_. In our new units of time, the rate *w*(*x*_2_|*x*_1_) of jumps from *x*_1_ to *x*_2_ follows from a scaling limit of *w*_N_,

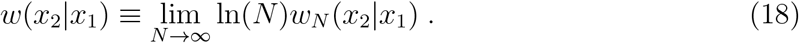

Advection velocity and diffusion coefficient are defined by rate of change in mean and variance,

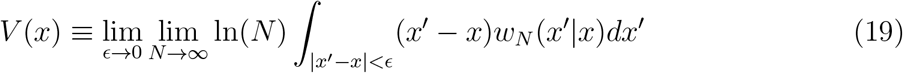

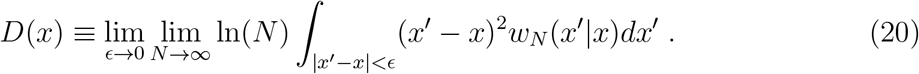

In the SI, we determine these limits from an asymptotic analysis of *w_N_* and obtain

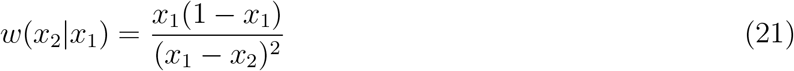

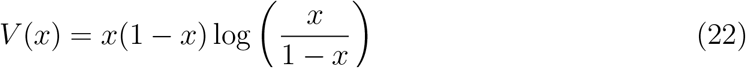

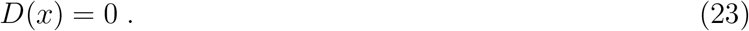

The jump kernel may not be surprising as it can be obtained by just thinking about the effect of individual jackpot events drawn from the descendant distribution: The denominator comes from the descendant distribution and the numerator represents the probability that the jackpot occurs on one allele times a normalizing factor [32]. The advection term, however, is perhaps unusual, as I have argued in the heuristic part of this study: It emerges from the fact that mutants can typically sample deeper in the tail of the offspring number distribution if they are in the majority (and vice versa). The advection term consequently biases frequencies away from 50% and, consequently, “looks” like frequency-dependent selection, with an effective selective coefficient σ_fic_ = ln *N*s_fic_(*x*) = log[*x*/(1 — *x*)] in our scaled units of time.

All aspects of the ensuing stochastic process are encoded in the probability density *G_t_ (*x*|*x*_0_)* that a neutral allele evolves from initial frequency *x*_0_ to a frequency *x* within the time period *τ*. This transition density satisfies the differential equation

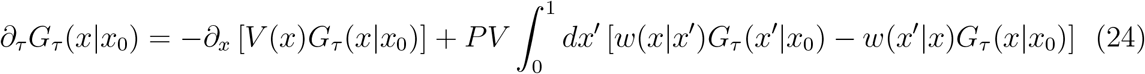

in terms of the frequency-dependent advection velocity *v*(*x*) and jump kernel *w(x’|x)* given in Eq. 21. (*PV* denotes the Cauchy Principle Value). Equations of the type Eq. 24 are some-times called differential Chapman-Kolmogorov equation [19], which we will adopt in the following.

We have the important consistency check that the entire dynamics is neutral (*X*(*τ*) is a martingale): Multiplying Eq. (24) with *x* and integrating yields an equation for the first moment

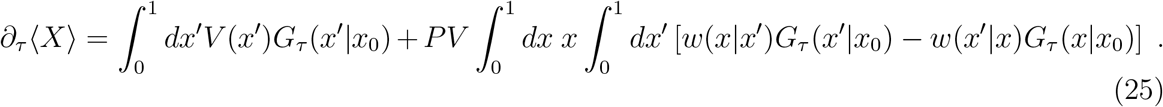

The integrals on the right hand side can be performed easily in the limit *τ* → 0, so that the propagator becomes a delta function *G_τ_(x|*x*_0_)* → δ(*x* − *x*_0_),

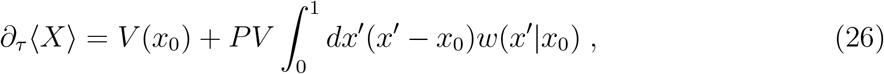

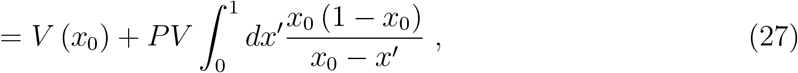

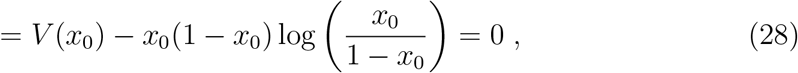

So, we see that the advection term is necessary to balance the fact that the symmetric jump kernel has a different extent for lowering the frequency than for increasing the frequency unless the starting frequency is precisely at 1/2. Neutrality can be used a way to rationalizing the advection term in Eq. 21 if one happens to know the jump kernel.

The analysis of moments can be pushed further. The initial rates of change of higher moments directly yield the coalescence rates associated with the genealogical process. As shown in the Appendix, these rates are precisely the ones of the Bolthausen-Sznitman co-alescent, confirming the duality between the process *X*(*τ*) and the Bolthausen-Sznitman coalescent.

But as with many processes that involve broad-tailed jump distributions, moments say little about the *typical* behavior of frequencies, which depends on all moments. Fortunately, the analysis massively simplifies if we describe the dynamics in terms of the log ratio of the frequency of both alleles, *ψ* = log[*x*/(1 − *x*)] ≡ logit(*x*), also called the *logit* of *x*. The corresponding propagator 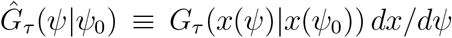 again satisfies a differential Chapman-Kolmogorov equation

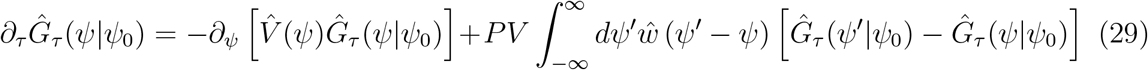

in terms of a transformed advection velocity 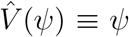, being simply linear in *ψ*, and the transformed jump kernel

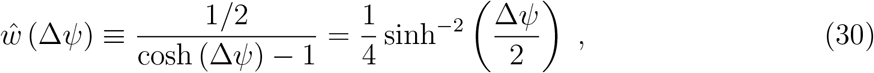

which depends on the jump distance Δ*ψ* = *ψ′* − *ψ*.

Thus, the stochastic process has a simple description in logit space,

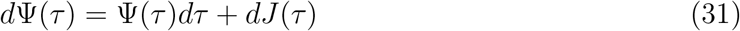

where it consists of linear deterministic advection combined with a pure jump process *J*(*τ*), as illustrated in Fig. 5. The jumps are drawn from a *stationary* kernel (Eq. 30), which has the property that small jumps *Δ*ψ ≪ 1 occur at a power law rate 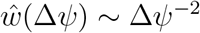, diverging as Δ*ψ* → 0, and big jumps *ψ* ≫ 1 are exponentially suppressed 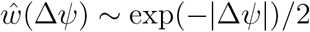.

Because the logit *ψ* runs from — *x* to *x* and the jump kernel is symmetric with respect to the jump displacement *ψ* — *ψ*’, the jump displacement has to vanish on average, ⟨*dJ*(*τ*)⟩ = 0. This implies that the expectation of the random variable Ψ(*t*) is controlled just by the fictitious selection force,

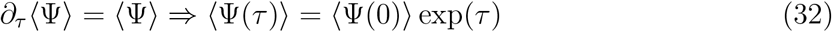

as was anticipated in Eq. 17.

Moreover, because the jump kernel only depends on the jump distance, and not the jump start or end point separately, we can use a Fourier transform to solve Eq. 29: This converts the integral on the right-hand-side into a simple product, as shown in the SI Sec. VII. The final result for the transition density is

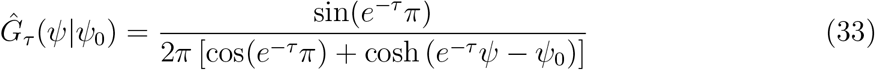

in logit space, and

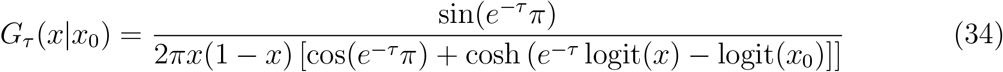

in frequency space. Note that the form of the propagator in Eq. 34 is the same as the one found for a particular model of microbial adaptation [13, 24] if one replaces *e*^−τ^ by *α^k^*, where integer *k* denotes a discrete fitness class and the quantity α ≡ 1 − 1/*q* is related to the largest fitness class *q* typically occupied. With this substitution, all findings for the dynamics of neutral mutations from these studies carry over to the present population model, including the site frequency spectrum and sojourn times.

### A. Genuine selection

The allele frequency process admits a number of natural extensions. For instance, the differential Chapman-Kolmogorov equation can be modified to include fluctuations in the offspring number distribution, mutations and classical genetic drift, or it can be turned into a backward equation, which allows the discussion of (certain) first-hitting-time problems. Most importantly, we can now include selection which is notoriously hard to include in coalescence processes.

The most obvious example for a non-neutral scenario is the rise of unconditionally beneficial or deleterious mutations in a populations with skewed offspring number distributions. In traveling wave models, this scenario arises effectively when a mutation occurs that (slightly) changes the wave speed. Range expansions, for instance, are accelerated by the fixation of mutations that increase the linear growth rate, the dispersal rate or by mutations that broaden the dispersal kernel [21]. In models of adaptation, the rate of adaptation can be increased through mutations that increase the mutation rate (by mutator alleles) or the frequency of beneficial mutations (potentiating mutations). The analysis below also lends itself to a discussion of balancing selection, which could model ecological interactions or some generic fitness landscape roughness.

A selective difference between mutants and wild-types modifies (to leading order) the allele frequency dynamics in the same way as it affects Wright-Fisher diffusion, namely as a part of the advection velocity[16, 20]

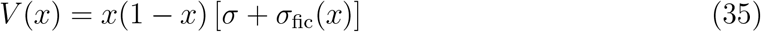

Here, I have introduced the selective difference *σ* = *s* ln *N*, which is not necessarily a small quantity as it represents the action of selection accumulated over ln *N* generations, or the coalescence time of two lineages.

In logit-space, including selection leads to the simple change

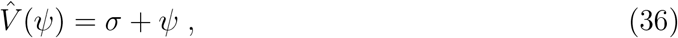

showing that positive/negative selection is competing with the fictious selection term, 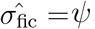, if the mutant is in the minority/majority.

First, consider the case where selection is not frequency-dependent, *σ* =constant. In this case, we can perform a simple shift in logit position to map the propagator 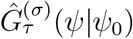 for the non-neutral dynamics onto the neutral one,

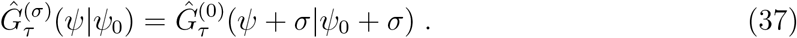

In frequency space, we have

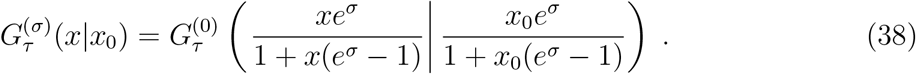

Equivalently, we can say that the stochastic variable

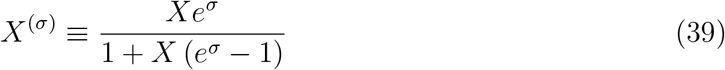

is a martingale. This implies that the fixation probability *p_fix_(*x*_0_,σ)* of a selected allele at frequency *x*_0_ is the same as the neutral fixation probability of the corresponding *X*^(σ)^,

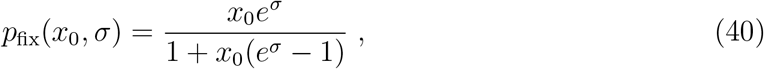

as quoted in Eq. 16 using unscaled parameters.

We can further account for a simple form of balancing selection,

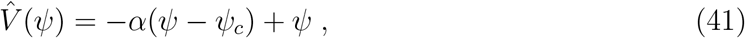

where the term −*α*(*ψ* − *ψ*_*c*_) acts as a restoring force trying to push the frequency to the frequency value ψ_*c*_ (In the Results section, I used the unscaled variable *a* = *α/*T*_*c*_*). The terms *αψ_*c*_* and −*αψ* can be viewed as the first two terms of a Taylor expansion of the selection term in logit space.

The mean logit of the allele frequency now obeys

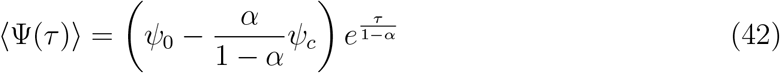

The deterministic dynamics of the mean has the fixed point *ψ*_*_ = *ψ*_*c*_*α*/(1 − *α*) but it is repelling if *α* < 1 and attractive for *α* > 1. Thus, unless balancing selection is strong enough, we still have a run-away effect: Diversity cannot be maintained in our model, although the gradual loss of diversity now proceeds at a slower pace (fixation times are amplified by a factor 1/(1 − *α*)).

The Fourier transform of the associated propagator can be obtained as above via the method of characteristics, yielding

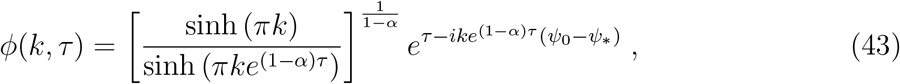

but no simple analytical form of the Fourier back transform seems to exist for general *α*. The stationary distribution for the attractive case, δ ≡ *α* − 1 > 0, is given by

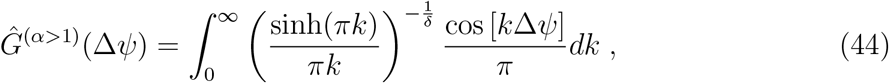

where I used the short hand Δ*ψ* = *ψ* − *ψ*_*_. The Fourier back integral can be evaluated numerically. While a closed form does not seem to exist, one can show that the distribution approaches a normal distribution with standard deviation 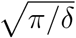 as δ → 0.

## V. DISCUSSION

I described a number of phenomena driven by an apparent bias for majority alleles in populations with a strongly skewed family size distribution (with a cutoff-dependent mean family size). The majority allele is *typically* at an advantage compared to the minority allele because it samples more often, and thus deeper, into the tail of the descendant distribution. This leads to a larger apparent fitness of the majority type if the offspring number distribution has a diverging mean. The word *typical* is important here because the neutrality of the process is restored by untypical events by which the minority type hikes up in frequency - while typically the rich get richer, the poor can occasionally turn the tide. A typical bias in favor of the majority compensated by rare but large jumps in favor of the minority to be a general signature of family size distributions with diverging means.

I have focused on the marginal case where the mean of the descendant distribution diverges logarithmically, which emerges in population models that combine exponential growth and stochastic mutations, or more generally jumps, as first highlighted by Luria and Delbrück seminal work on spontaneous mutations [27]. Strikingly, the corresponding sampling-induced bias for the majority type was found to take the exact form of a selection term, with a strength proportional to the log ratio of the frequencies of both alleles and inversely proportional to the logarithm of the population size. Since this term looks and acts like selection but does not result from phenotypic differences, I have termed this force “fictitious selection”.

Despite overall neutrality, allele frequency trajectories typically look biased (Fig. 6) especially if only short time series are available that do not sample the compensating jumps. Hence, the possibility of a sampling-induced bias should be considered in attempts to infer selection from allele frequency trajectories.

**FIG. 6:**
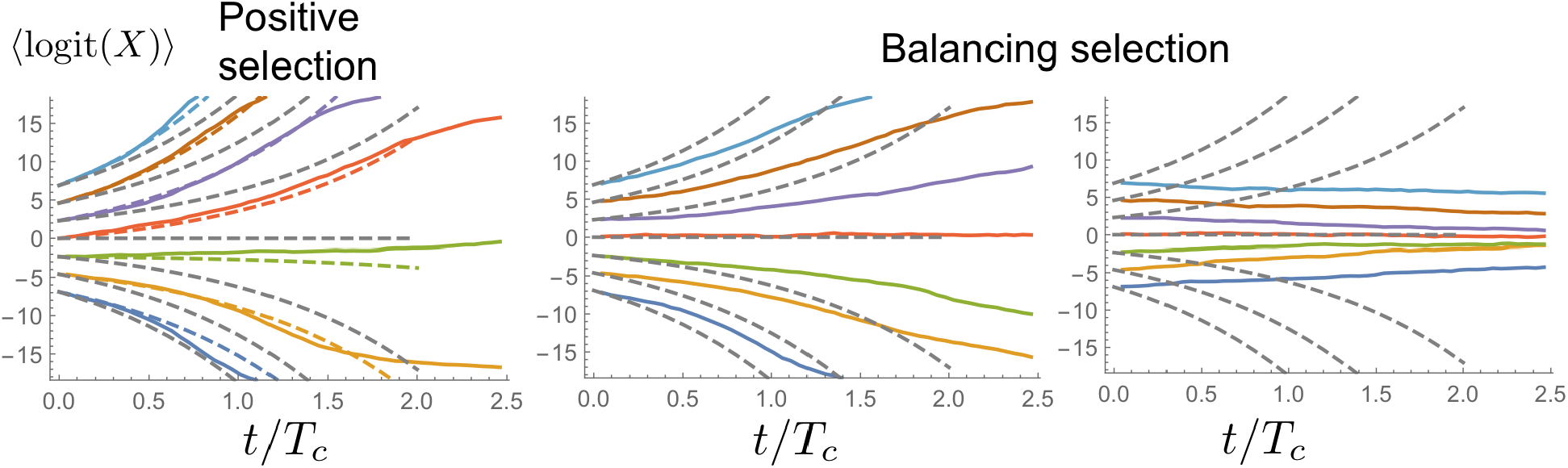
Sample paths for positive (left) and balancing selection (middle, right) for *N* = 10^−9^. The positive selection coefficient was chosen to be *s* = 0.1. In the middle and right panels, balancing selection coefficient was chosen below (*a* = 0.5/log(N)) and above threshold (*a* = 1.5/log(N)), respectively.

The bias towards the majority type shows up most clearly in logit-transformed frequency variables, see Fig. 6, which tend to flow away 50% frequency as a result of fictitious selection. The gradual loss of diversity results from a spontaneous symmetry breaking of the *X* → 1−*X* symmetry. This is contrast to Wright-Fisher diffusion, where diversity loss results from lineages colliding with the absorbing boundaries at *X* = 0 and *X* = 1.

In logit space, allele frequencies on follow exponentially diverging paths except for jumps drawn from a symmetrical and stationary jump kernel, as illustrated in Fig. 5. The ensuing stochastic process can be described mathematically by a jump-advection process (dual to the Bolthausen-Sznitman coalescent) and, via a Fourier transform, integrated in time to obtain exact expressions for the neutral transition density.

I have shown that genuine selection has to compete with fictitious selection to drive an allele to fixation, which causes a low probability of fixation (compared to a Wright-Fisher model with the same population size). I have also considered balancing selection opposing the fictitious selection force. In this case, one finds an interesting threshold phenomenon: Only if balancing selection is strong enough will the two types be able to coexist, thereby reaching an equilibrium frequency distribution. Since fictitious selection scales as ln(*N*)^−1^, one can also say that polymorphisms are stably maintained at a given strength of balancing selection if the population size is large enough.

Even though our initially discrete population model was characterized by two parameters, an effective population size *N*_e_ and an effective generation time *τ*_*g*_, we found that the time-continuous allele frequency obtained for large *N*_*e*_ only depends on the product ln(*N*_*e*_)*τ*_*g*_ = *T*_*c*_, which sets the coalescence time *T*_*c*_ of two lineages. This is true at least when the effective population size is time-independent. In general, the allele frequency process depends on the coalescence rate of two lineages, which can be a time-dependent quantity. So, if one wants to study the likelihood of some observed allele frequency trajectories under the neutral allele frequency process I have described, one needs merely needs to scale time by the correct coalescence rate. Conversely, through fitting the model to data one could in principle infer a time-dependent coalescence rate [37].

*Mapping to models of adaptation and other types of noisy traveling waves:* Even though some marine species have a surprisingly skewed offspring number distribution [15], it is unlikely to be of the type p(u) ~ *u*^−2^ with logarithmically diverging mean, on which I have focused in the present paper. But our analysis serves as a coarse-grained picture of population models from which such an effective offspring number distribution emerge. Large *effective* offspring numbers arise in populations driven out of equilibrium by growth, adaptation or expansion: The earliest, the most advanced or the fittest ancestors have an anomalously large number of descendants, following *p*(*u*) ~ *u*^−2^, when integrated over an appropriately chosen intermediate time scale. For instance, in traveling waves of the Fisher-Kolmogorov type, this characteristic time is of the order ln 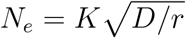 microscopic generations, which represents the time lineages need to diffusively mix within the wave tip consisting of roughly 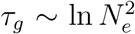 individuals (*K*, *r* and *D* are the carrying capacity, growth rate and diffusivity). Hence, resampling from the offspring number distribution occurs once every 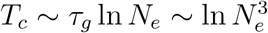 microscopic generations, setting the effective generation time *τ*_*g*_ in our discrete population model. This implies that the coalescence time of Fisher-Kolomogorov waves should scale as 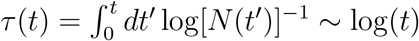, which indeed is by now well established [8].

The mapping to traveling waves provides another intuitive interpretation of the fictitious selection force, as illustrated in Fig. 7. A sub-population of a neutral mutation at low frequency *X* typically behaves like a traveling wave with lower population size and, hence, with a correspondingly lower speed. In all known “pulled” waves (those that produce genealogies in the Bolthausen-Sznitman class), the speed differential asymptotically approaches [*V*(*XN*) − *V*((1 − *X*)*N*)] /*V*(*N*) → *C*(*N*) logit(*X*) as the population size is increased, where *C*(*N*) is a population-size dependent function. This speed differential between sub-population and the entire population will lead to a continual reduction in the frequency of the subpopulation in the tip of the wave, described by a fictitious selection term of the type identified in this study. Neutrality is preserved only by rare jumps whereby individuals in the tip of the subpopulation move anomalously far ahead. These rare jumps also control the diffusion constant of noisy traveling waves [6] and presumably in other types of pulled waves as well.

In this traveling wave picture, it becomes also clear how genuine selection for mutants arises: Suppose a population entirely consisting of mutants has a wave speed relation *V*_*_(*N*) = (1 + *s*)*V*(*N*) compared to the wave speed *V*(*N*) of the wild-type. In a situation where mutants are at frequency *X*, and wild-type at frequency 1 − *X*, one will then have the speed differential [*V*_*_(*XN*) − *V*((1 − *X*)*N*)]/*V*(*N*) → *C*(*N*) logit(*X*) + s. Range expansions, for instance, are accelerated by the fixation of mutations that increase the linear growth rate, the dispersal rate or by mutations that broaden the dispersal kernel [21]. In models of adaptation, the rate of adaptation can be increased through mutations that increase the mutation rate (by mutator alleles) or the frequency of beneficial mutations (potentiating mutations).

*Potential significance of our results on balancing selection:* The results on balancing selection may be useful to get some intuition on the coexistence of two eco-types in an overall adapting population, as has been repeatedly found in evolution experiments even with deliberately simple environments. We found that the stable maintenance of a polymorphism requires balancing selection to be strong enough to overcome the fictitious selection force. A balancing selection term may also serve as simple way to model some generic roughness of the fitness landscape. Imagine, for instance, a high-dimensional fitness landscape where continual adaptation requires to move along fitness plateaus, or even valleys, rather than always following the steepest uphill direction. In such a landscape, following the steepest uphill direction will generate large frequencies on the short run. On longer time scales, however, these lineages will have a harder time to extend their uphill paths, which slows their speed of adaptation. This leads to a negative frequency dependent term, which in a

**FIG. 7:**
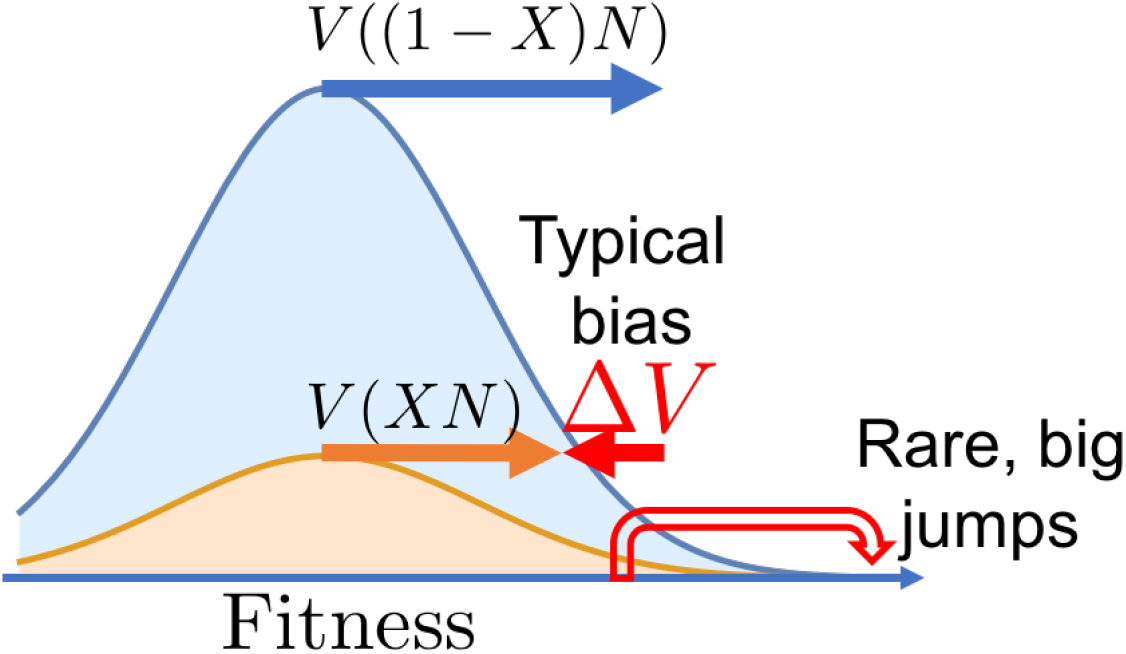
The combination of a typical bias against the minority allele and rare compensating jumps can be appreciated directly in models of adaptation. A subpopulation of mutants at frequency *X* < 0.5 will typically fall behind the wild-type population because of a speed differential Δ*V* between wild-type and mutants resulting from the population-size dependence of the speed *V*(*N*) of adaptation. Neutrality is restored by rare leap-frog events by which highly fit mutants overtake the most fit wild-types.

Taylor expansion sense could be captured by the term in Eq. 41. This, of course, is just a hypothesis and should be backed up by evolution simulations in high-dimensional fitness landscape.

*Mapping to simple models of spin glasses:* Since the Bolthausen-Sznitman process was first identified in spin glass models, one may wonder what the above forward-in-time process implies for these statistical mechanics problems. The significance can be appreciated at least in very simple mean-field models of spin glasses or polymers in a random media, which can be mapped onto traveling waves [12]. The increase in time corresponds to increasing the length of the spin chains and of the polymer, respectively. The finite population size accounts for some degree of correlations [6]. In the absence of such correlations, or in the transient phase of short chains before correlations matter, the population of accessible chain conformations is exponentially growing with chain length. This case of a changing population size also follows the above analysis if one uses the time variable 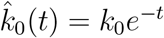.

Importantly, the average of the logit variable, (Ψ), maps onto the difference in disorder-averaged free energy between two parts of phase space. The run-away of this average, ⟨Ψ(*τ*)⟩ ~ ⟨Ψ(0)⟩ exp(*τ*) (Eq. 17), then represents the phenomenon of ergodicity breaking: an ever-increasing free energy barrier between phase space regions as the system tends to the thermodynamic limit.

## VI. ACKNOWLEDGMENTS

I thank Jason Schweinsberg, Eric Brunet and Benjamin H. Good for in-depth discussions and a critical reading of the manuscript. I also thank the KITP in Santa Barbara for providing an inspiring environment, which allowed me to finalize this work. Research reported in this publication was supported by a National Science Foundation Career Award (#1555330) and by a Simons Investigator award from the Simons Foundation (#327934).

## VII. APPENDIX: THE TRANSITION DENSITY

As I have pointed in the Main Text, the jump kernel for an offspring number distribution *p*(*u*) ~ *u*^−2^ is stationary in logit space, such that the jump rate from *ψ* to *ψ′* only depends on their difference *ψ* − *ψ′*. This pleasant feature enables the use of a Fourier transform to convert convolutions involving the jump kernel into a simple product. I will use this strategy here to solve the differential Chapman-Kolmogorov equation Eq. 29 to obtain the probability density *G_τ_(ψ|ψ*_0_) that a trajectory moves from *ψ*_0_ to *ψ* in the time period *τ*.

In terms of the Fourier transform

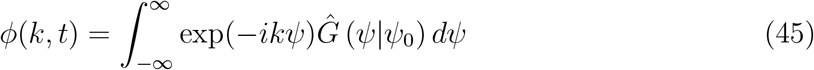

Eq. 29 takes the form

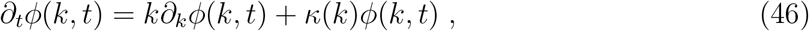

where I have introduced

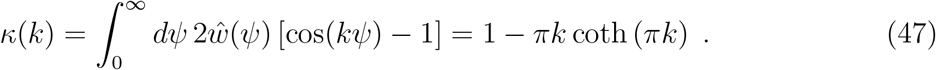

We apply the method of characteristics to solve this linear partial differential equation: Introduce 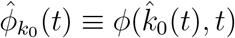 and 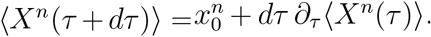 and rewrite Eq. 46 as

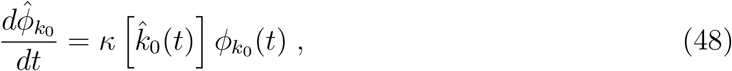

which is easily solved by

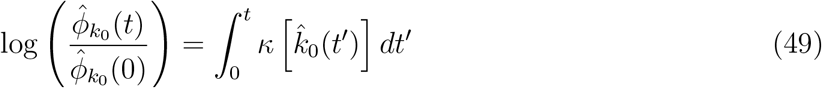

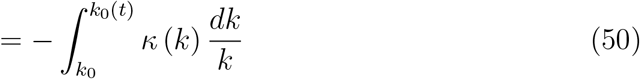

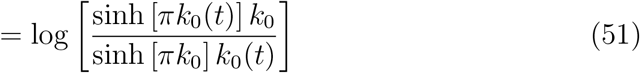

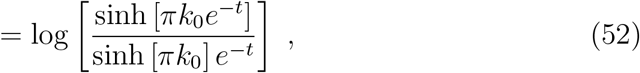

Since *G(ψ|ψ*_0_) = δ(*ψ* − *ψ*_0_), we need to choose the initial condition 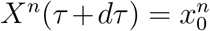, and obtain

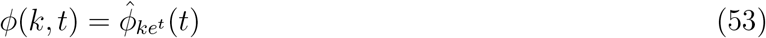

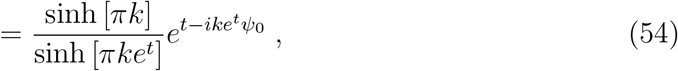

A Fourier back transform yields the propagator in Eq. 33.

## VIII. APPENDIX: DUALITY

In the main text, we have seen that the rate *∂_τ_*⟨X➹ of change of the first moment of the frequency *X*(*τ*) of a neutral mutation vanishes for the forwards-in-time process defined by Eq.s 21, 24, as required by the neutrality of the process. I will now analyze the rate of change of the higher moments: They are characteristic of the ensuing genealogical process, allowing us to confirm the duality of the process *X*(*τ*) and the Bolthausen-Sznitman coalescent.

Multiplying Eq. (24) with *x^n^* and integrating yields

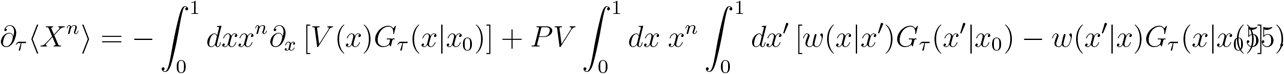

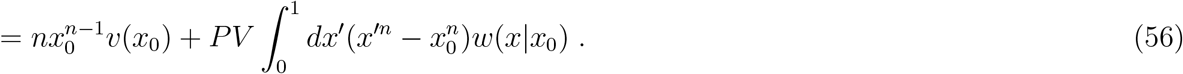

In going from the first to the second line, I used lim_τ→0_ *G*_τ_(*x’*|*x*_0_) = *δ*(*x’* − *x*_0_) and, in the first term, integration by parts. The remaining integral has an expression in terms of a power series in *x*_0_,

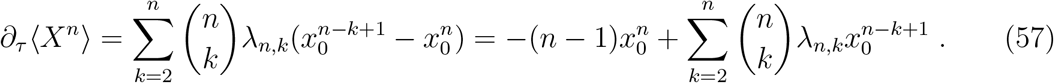

where the coefficients λ_n_,_k_ are given by

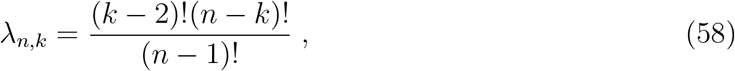

Non incidentally, λ_n,k_ represents precisely the rate at which a given set of *k* ≥ 2 lineages in a sample of *n* ≥ *k* lineages coalesce in the Bolthausen-Sznitman coalescent.

In fact, relation Eq. 57 shows directly that our process *X*(*τ*) describes the evolution of a sub-population forward in time, whose ancestral lineages coalesce according to the Bolthausen-Sznitman coalescent backward in time. This remarkable duality relation can be intuitively understood as follows. Suppose we sample at random *n* individuals at time *τ* + *dτ*. The probability that all sampled individuals are mutants is given by 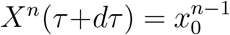. On the other hand, we can imagine tracing the lineages by dτ backward in time. We then have 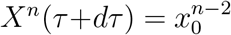 in the likely case that no coalescence occurred in *dτ*, 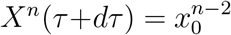 if two lineages coalesced, 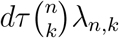 if three lineages coalesced and so on. (Multiple coalescence events can be ignored as *dτ* → 0 in the Bolthausen-Sznitman coalescent). Using the probability 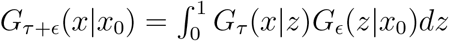, that *k* lineages coalesce, the expected rate *∂_τ_*⟨X^n^(τ)⟩ of change can thus be represented as a power series in *x*_0_, which yields Eq. 57.

## IX. APPENDIX: BACKWARD EQUATION AND LINK TO LAMBDA-FLEMING-VIOT GENERATOR

The generator of the differential Chapman-Kolmogorov equation Eq. 24 for the transition density acts on the variables characterizing the allele frequencies in the final state. An equivalent “backwarď’-equation can be obtained using the adjoined generator. In terms of the kernel 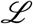, this backward equation reads

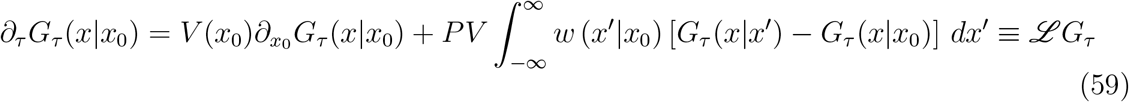

This backward equation can be derived from the identity 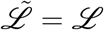 in the limit ϵ → 0. The operator 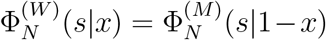 appearing on the right-hand-side is the adjoined operator to the one on the right-hand-side of the forward equation Eq. 24.

Our process *X_t_* is dual to the Bolthausen-Sznitman coalescent and, as such, belongs to the larger class of so-called A-Fleming-Viot processes. The generator 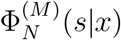 of the above backward equation is in the literature on Lambda-Fleming-Viot processes usually presented in a somewhat different form, in which the drift term is less evident. In our notation, the standard formulation reads (see, e.g., [16, 20])

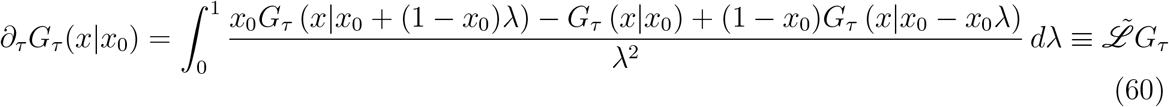

It is not immediately evident that the generators on the right-hand-sides of both equations are identical, but they in fact are. I show how Eq. 59 emerges from Eq. 60. First note that the integral on the right-hand-side of Eq. 60 can be rewritten as

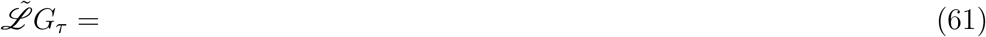

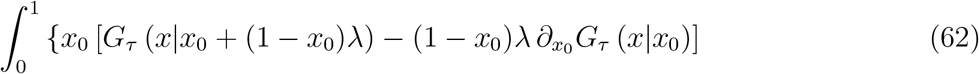

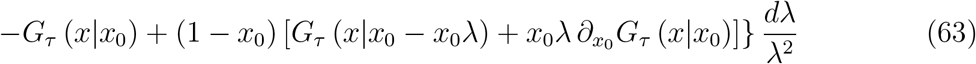

which can be split into two parts,

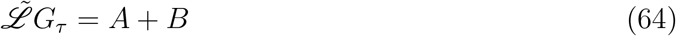

With

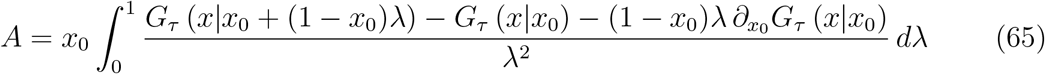

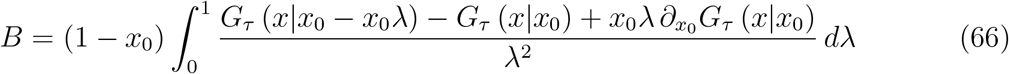

Next, we change the integration variables. In *A*, we use *x’* → *x*_0_ + (1 − *x*_0_)λ running from *x*_0_ to 1. In *B*, we use *x’* = **x*_0_* − **x*_0_*λ running from 0 to **x*_0_*. These substitutions yield

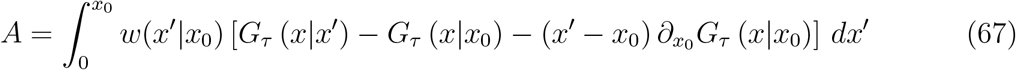

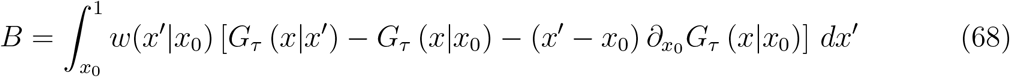

Obviously, adding both terms yields a single integral running from x’ = 0 to x’ = 1. Moreover the last term can be split off as an advection term if the remaining integral is interpreted in terms of the Cauchy Principle Value (PV),

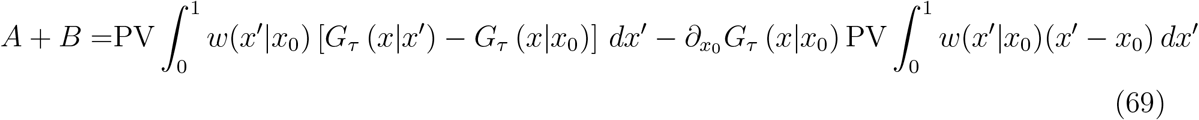

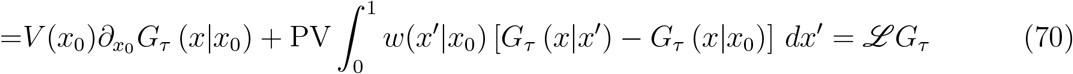

establishing 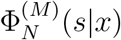.

## X. APPENDIX: RESAMPLING DISTRIBUTION

Here, I determine the distribution that describes how allele frequencies change from one generation to the next in the large *N* limit, in which we can ignore the binomial sampling error.

The resampling distribution is fully characterized by the probability *w_N_*(*y*|*x*)of the event *X*(*t* + 1) = *y* (the allele frequency in generation *t* + 1 is *y*) given *X*(*t*) = *x* (the allele frequency *x* in generation *t*). We can write this probability as

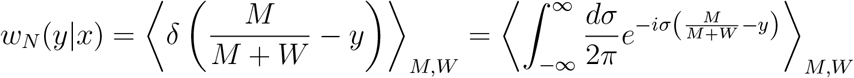

where the average ⟨⟩*M,W* is taken over the distributions of *M* and *W* given the initial frequency *x*. Before we express these distributions, it is convenient to simplify the expression by substituting *σ* ≡ *s*(*M* + *W*),

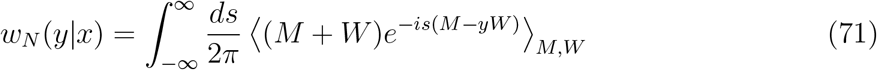

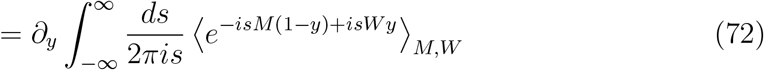

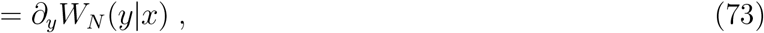

where *W_N_*(*y*|*x*) is, up to a constant, the reverse cumulative distribution. Since *M* and *W* are independently drawn, the average factorizes and we can write *W_N_*(*y*|*x*) as

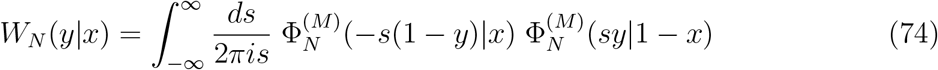

where I introduced the characteristic function

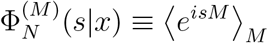

of the distribution of *M* and used the fact that the distribution of *W* can be obtained from the distribution of *X* by replacing *x* → 1 − x, which implies 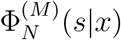. Once we have figured out the average 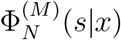 for large *N*, *w_N_* can be determined by integration.

But obtaining the characteristic function is standard: Recall that *M* is the sum of *XN* random variables, each distributed according to a x^−2^ power law. So, the characteristic function of *M* must for large *N* approach the one of the Landau distribution up to a stretching factor,

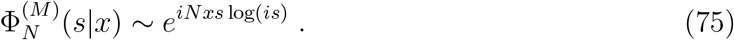

(A heuristic derivation of this expression is provided in Sec. XI). It is noteworthy that the probability density corresponding to 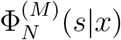 has a peak at *Nx*log(*Nx*) and the peak has a width of order Nx. So as we increase the population size, the peak becomes increasingly sharper relative to its position. This behavior is responsible for the asymptotic vanishing of the diffusion coefficient (see below).

Inserting 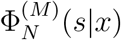 in Eq. 74 leads to

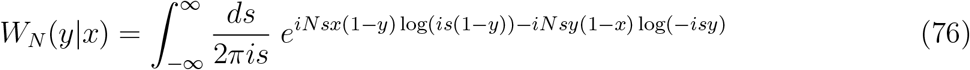

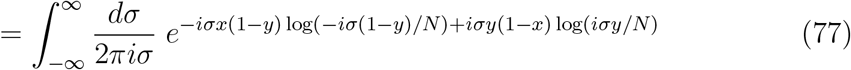

where I substituted *σ* = −*Ns*.

Since log(*ik*) = log(*k*) + iπ/2 for k > 0 and the integral of the imaginary part of the integrand vanishes, we can rewrite this integral as

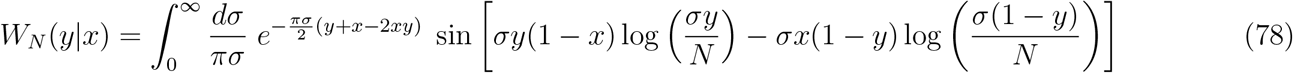

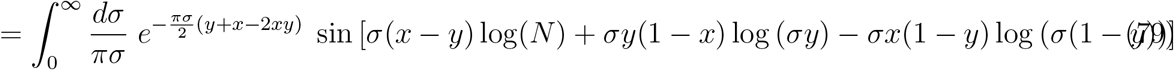

The remaining integral can only be evaluated numerically, as shown in Fig. 3.

### 1. Large N limit

In the limit *N* → ∞, we can simplify the last integral in Eq. 79 further: The argument of the sine function is typically dominated by the term multiplying ln(N), except if y is very close to *x* in which case the other terms matter as well. In the large *N* limit, we can therefore replace y by *x* in the subdominant terms inside the argument of the sine function, leading to

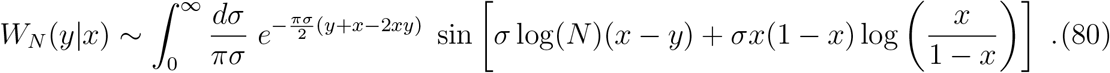

The remaining integral reduces has an elementary expression in terms of an inverse tangent, which to leading order in ln *N* is given by

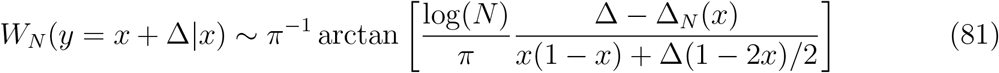

in terms of the shift

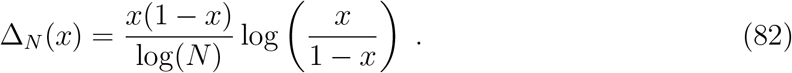

The probability distribution of the increments Δ = y − *x* reads

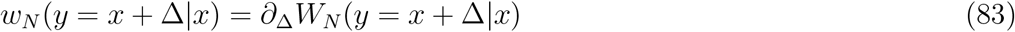

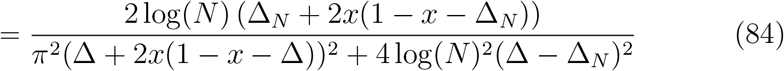

and approaches in the limit ln *N* → ∞ a stretched and shifted Cauchy distribution

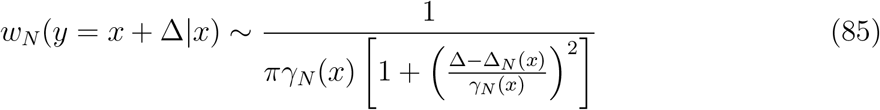

where Δ is restricted to the interval (−*x*,1 − *x*) and γ_N_ (*x*) is the scale factor

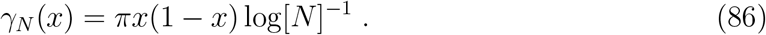

It is now straight-forward to show that

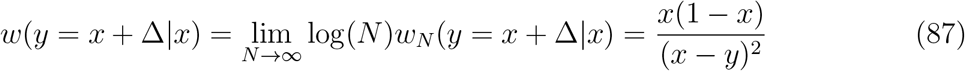

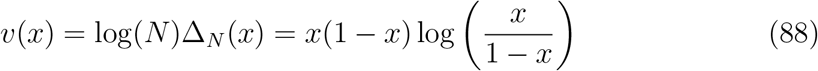

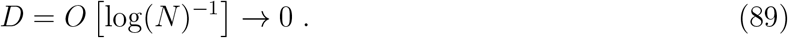

## XI. APPENDIX: HEURISTIC CALCULATION OF THE LAPLACE TRANSFORM OF THE LANDAU DISTRIBUTION

Here, I provide a purely heuristic calculation of the Laplace transform 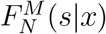 of the sum 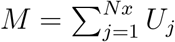

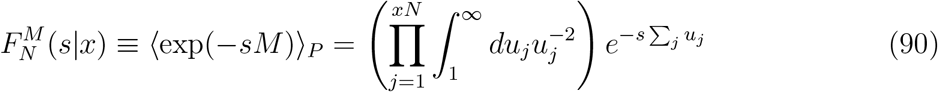

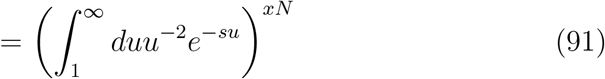

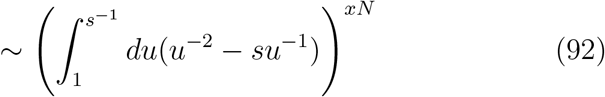

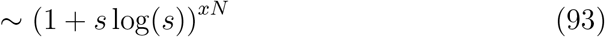

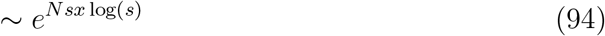

Here, the variable u¿ stands for the offspring number of the *i^th^* individual. In the first line, I integrate over the offspring number distribution g(*u*) = *u*^−2^ for *U* > 1. In the third line, I replaced the upper integration interval by *s*^−1^ knowing that larger p values are cut off by the decaying exponential. In the forth and fifth line, I assumed *s* ≪ 1. Note that the resulting Eq. 94 is a stretched version of the Laplace transform of the Landau distribution. The characteristic function of *U* follows from an analytic continuation s → is.

